# Genome-wide identification of lineage and locus specific variation associated with pneumococcal carriage duration

**DOI:** 10.1101/107086

**Authors:** John A. Lees, Nicholas J. Croucher, Goldblatt David, Nosten Francois, Parkhill Julian, Turner Claudia, Turner Paul, D. Bentley Stephen

## Abstract

*Streptococcus pneumoniae* is a leading cause of invasive disease in infants, especially in low-income settings. Asymptomatic carriage in the nasopharynx is a prerequisite for disease, and the duration of carriage is an important consideration in modelling transmission dynamics and vaccine response. Existing studies of carriage duration variability are based at the serotype level only, and do not probe variation within lineages or fully quantify interactions with other environmental factors.

Here we developed a model to calculate the duration of carriage episodes from longitudinal swab data. By combining these results with whole genome sequence data we estimate that pneumococcal genomic variation accounted for 63% of the phenotype variation, whereas host traits accounted for less than 5%. We further partitioned this heritability into both lineage and locus effects, and quantified the amount attributable to the largest sources of variation in carriage duration: serotype (17%), drug-resistance (9%) and other significant locus effects (7%). For the locus effects, a genome-wide association study identified 16 loci which may have an effect on carriage duration independent of serotype. Hits at a genome-wide level of significance were to prophage sequences, suggesting infection by such viruses substantially affects carriage duration.

These results show that both serotype and non-serotype specific effects alter carriage duration in infants and young children and are more important than other environmental factors such as host genetics. This has implications for models of pneumococcal competition and antibiotic resistance, and leads the way for the analysis of heritability of complex bacterial traits.

**Significance statement:** Other than serotype, the genetic determinants of pneumococcal carriage duration are unknown. In this study we used longitudinal sampling to measure the duration of carriage in infants, and searched for any associated variation in the pan-genome. While we found that the pathogen genome explains most of the variability in duration, serotype did not fully account for this. Recent theoretical work has proposed the existence of alleles which alter carriage duration to explain the puzzle of continued coexistence of antibiotic-resistant and sensitive strains. Here we have shown that these alleles do exist in a natural population, and also identified candidates for the loci which fulfil this role. Together these findings have implications for future modelling of pneumococcal epidemiology and resistance.

## Introduction

*Streptococcus pneumoniae* is a human pathogen that can cause diseases such as pneumonia, otitis media and meningitis. Pneumococcal disease burden is highest in children (1). For disease to be caused pneumococci must first transmit to the host, colonise the nasopharynx and finally cross into an invasive niche. The pneumococcus spends most of the transmission cycle in the nasopharynx, and so understanding and predicting the amount of time spent in this niche is critical for understanding this bacterium’s epidemiology, and therefore controlling transmission (2, 3).

The nasopharynx is a complex niche in which each pneumococcal genotype must tackle a wide range of factors including host immune defence (4), other bacterial species (5), and other pneumococcal lineages (6, 7) in order to maintain the genotype’s population. The average nasopharyngeal duration period is therefore affected by a large number of factors, which may, themselves, interact.

One factor that is known to strongly associate with carriage duration is serotype: as capsular polysaccharides are important in bacterial physiology and determining host immune response, different serotypes have different clearance and acquisition rates (3, 8–11). Additionally, a range of other proteins have been identified as critical to the colonisation process (12), some of which exhibit similar levels of diversity to the capsule polysaccharide synthesis locus (13, 14). However, the overall and relative contributions of these sequence variations to carriage rate have not yet been characterised. In addition variation of pathogen protein sequence, accessory genes and interaction effects between genetic elements may also have as yet unknown effects on carriage duration.

Changes in average carriage duration have been shown to be linked with recombination rate (15), which has been found to correlate with antibiotic resistance (16) and invasive potential (15). The carriage duration by different serotypes is widely used in models of pneumococcal epidemiology, and consequently is important in evaluating the efficacy of the pneumococcal conjugate vaccine (PCV) (2, 17). Additionally, modelling work has proposed that if alleles exist which alter carriage duration, these explain the coexistence of antibiotic-resistant and sensitive strains in the population (18). Measurement of carriage duration and the analysis of its variance beyond the resolution of serotype will have important consequences for these models.

We sought to determine the overall importance of the pathogen genotype in carriage duration in a human population, and to identify and quantify the elements of the genome responsible for the variation in carriage duration. By combining epidemiological modelling of longitudinal swab data with and genome wide association study methods on the connected sequences, we made heritability estimates for carriage duration. We further partitioned the heritability into contributions from lineage and locus effects (19) to quantify the variation caused by each individual factor.

## Results

### Ascertainment of carriage episode duration using epidemiological modelling

We first estimated carriage duration from longitudinal swab data available for the study population. For 598 unvaccinated children a complete set of 24 swabs taken over a two year period were available, an extension on the previous study (11, 20). We only considered swabs from infants in the study, as mothers did not have sufficient sampling resolution relative to their average length of carriage to determine carriage duration. Furthermore, the immune response of mothers to bacterial pathogens is different to children (21), leading to shorter carriage durations (22).

To estimate carriage duration from the longitudinal swab data we constructed a set of hidden Markov models (HMMs) with hidden states corresponding to whether a child was carrying a serotype at a given time point, and observed states corresponding to whether a positive swab was observed for this serotype at this time point.

The most general model for the swab data would be a vector with an entry of 0 or 1 for every possible serotype (of 56 observed in the population), corresponding to whether each serotype was observed in the swab at each time point. However, the number of parameters to estimate in this model (with over 6 million states) is much larger than the number of data points (around 14000), and in particular some serotypes have very few positive observations. Instead, we modelled each serotype separately.

The models fitted, and their permitted transitions and emissions are shown in figure 2. In model one, observation *i* emits state 2 if positively swabbed for the serotype, and state 1 otherwise. The unobserved states correspond to the child ‘carrying’ and being ‘clear’ of the serotype respectively. We assume swabs have a specificity of one, so do not show positive culture when the child is clear of the carried serotype; we therefore set the coefficient for the chance of observing positive culture when no bacteria are present to zero (*e*_12_ = 0 in the emission matrix). Model two adds a third state of ‘multiple carriage’ which is occupied when the serotype and at least one other are being carried. Both models were compared with a version which allows the parameters to covary with whether the child has carried pneumococcus previously. Model three accounts for this explicitly by having separate states and emissions based on whether carriage has previously been observed.

**Figure 1:**
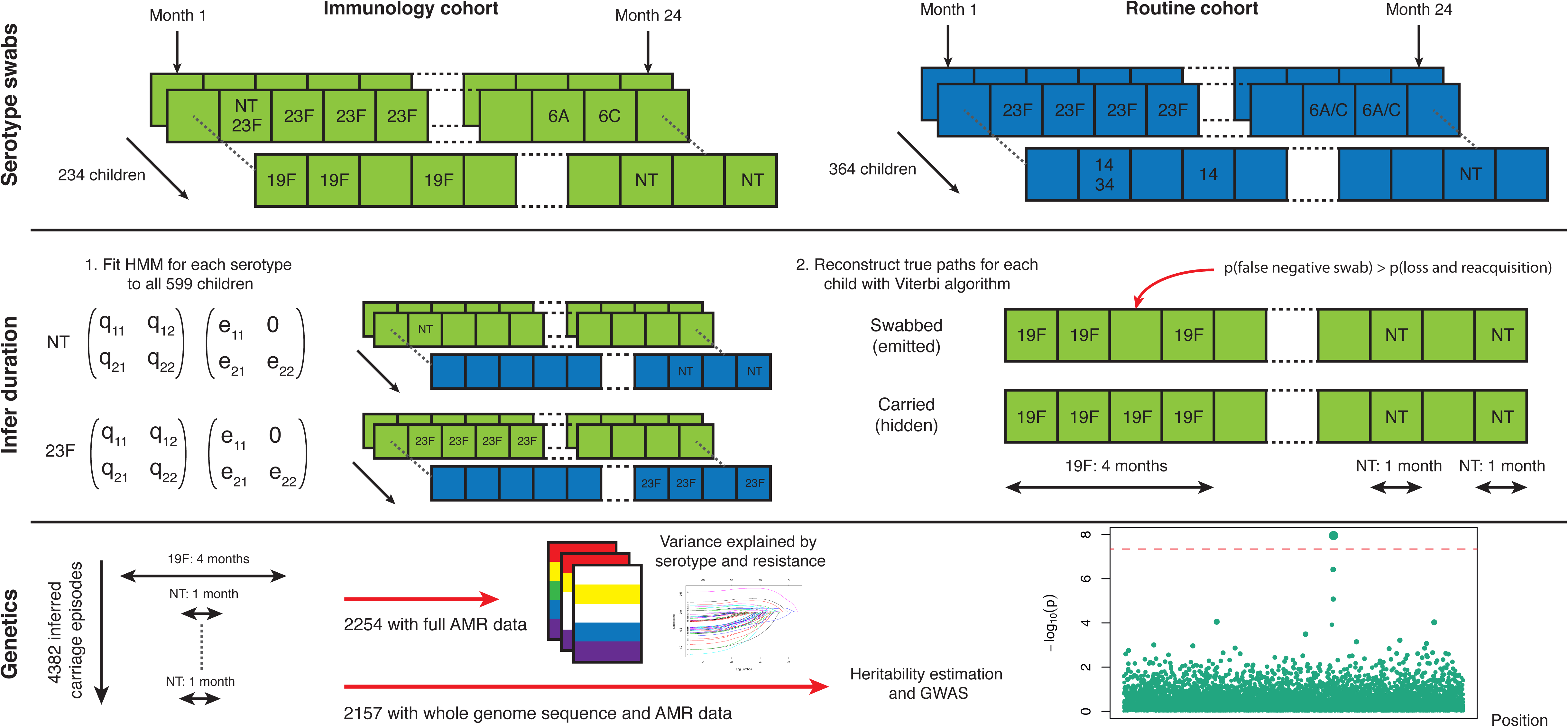
Swabbing and sequencing study design. We start with serotype swab data on 598 children from two cohorts, taken every month after birth for two years. For all samples we fitted the transition and emission probabilities of a continuous time hidden Markov model for each serotype. Then, for each child, we used these parameters were then used to infer the most likely carriage durations. We matched carriage episodes with resistance and genomic data to draw conclusions on the basis of variation in this epidemiological parameter.

**Figure 2:**
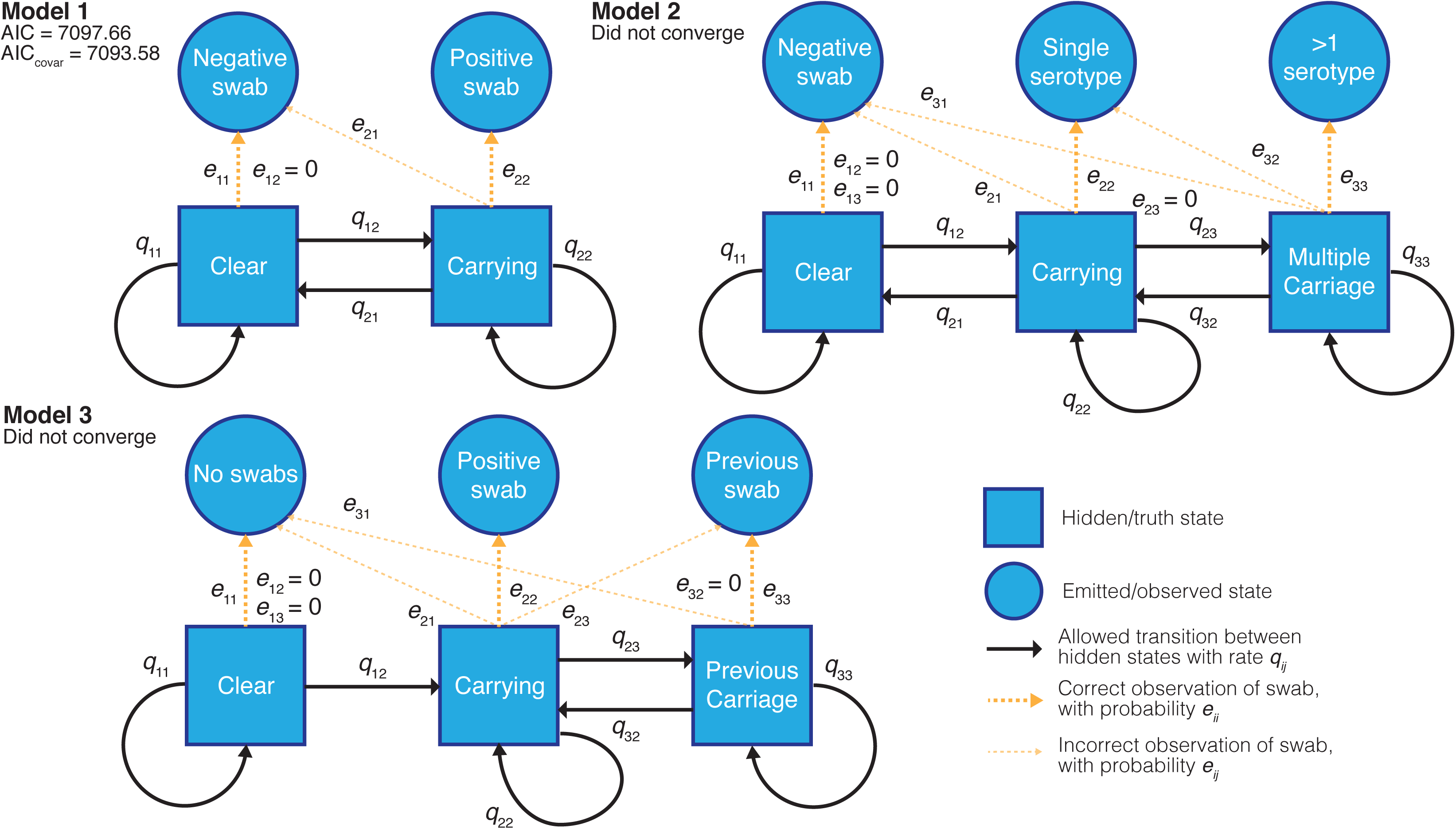
Hidden Markov models of swab time series, and their goodness-of-fit. We fitted three different models to the processed time-series data with states, allowed transitions and emissions as shown. We refitted each model allowing the transitions probabilities to covary with the age of the child and whether the child had carried pneumococcus previously. For the converged model the Akaike information criterion (AIC) is shown for the original fit, and when including these covariates (AIC_covar_).

We applied all the models to 19F carriage episodes, as these had the most data available, and calculated the Akaike information criterion (23) for each model that converged. We then fitted the best performing model in this test for all serotypes separately. 6A and 6C were treated as a single serotype, as they were not always distinguished in the course of the study. The models for 19F, 23F, 6A/C, 6B, 14 and non-typable (NT) converged, but other serotypes did not have enough observations to successfully fit the parameters of the model. For these less prevalent serotypes we used the transition and emission parameters from the 19F model fitted with the correct observations when reconstructing the most likely route taken through the hidden states. Results were inspected to ensure this did not cause systematic overestimation when compared with previous studies.

We found that the fit for NT swabs produced results which overestimated carriage duration when compared to previously reported estimates. The best fit to the model overestimated the *e*_21_ parameter, which measures the false negative rate of swabbing, in favour of reduced transition intensities. We therefore fitted the model again, fixing this rate at 0.12. We based this figure on non-typable *Streptococcus pneumoniae* abundance as defined by 16S survey sequencing. At 1% proportional abundance in the sample, 12% came out as culture negative (SI table 1).

From all the swab data, we estimated that there were a total of 4382 carriage episodes, of which 2254 had a complete set of AMR data available. After removing ten outlier observations from accidentally taken swabs (SI figure 1), we were able to match 2157 sequenced genomes with a carriage duration. Duration was positively skewed due to some observations of very long carriage times. We therefore took a monotonic transform of the carriage duration to maximise the study’s power to discover associations and estimate heritability (SI figure 2).

### Overall heritability of carriage duration is high

The variation in carriage duration 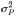 is partly caused by variance in pneumococcal genetics, and variance in other potentially unknown factors such as host age and host genetics. It is common to write this sum as two components: genetic effects 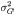 and environmental effects 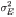. The proportion of the overall variation which can be explained by the genetics of the bacterium is known as the broad-sense heritability 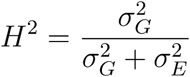. Variants which are directly associated with carriage duration independently of other variants (non-epistatic effects) contribute to the narrow-sense heritability *h*^*2*^, which is smaller than the overall broad-sense heritability (24).

*H*^2^ can be estimated by linear regression on the phenotype of donor-recipient pairs which nearly share their genetics (25). However in this dataset we were only able to confidently identify five transmission events, which was not enough to apply this method. Alternatively, analysis of variance of the phenotype between pathogens with similar genetics can be used to estimate heritability (26). By applying this to phylogenetically similar bacteria, we estimated that *H*^2^ = 0.634 (95% CI 0.592-0.686). This implies that the genetics of *S. pneumoniae* is the most important factor in determining carriage duration.

A lower bound on *h*^2^ can be calculated by fitting a linear mixed model through maximum likelihood to common SNPs (*h*^2^_*SNP*_) (27, 28). We used the model in warped-lmm (29) to estimate (*h*^2^_*SNP*_) for carriage duration data, yielding an estimate of 0.445, consistent with our estimate for *H*^2^.

### Serotype and drug resistance explain part of the narrow-sense heritability

After calculating the overall heritability, we wished to determine the amount that the specific variation in the pathogen genome contributes to changing carriage duration. In the context of genome wide association studies (GWAS) in bacteria strong linkage-disequilibrium (LD) is present across the entire genome, making it difficult to pinpoint variants associated with carriage duration and not just present in the background of longer or shorter carried lineages (30). In *S. pneumoniae*, serotype and antibiogram are correlated with the overall genome sequence (31–33). If these factors are associated with carriage duration, large sets of variants which define long-carried and short-carried lineages will be correlated with carriage duration in a naive association test (30, 34).

A distinction has therefore been made between variants which evolve convergently and affect a phenotype independently of lineage – termed locus effects – to those which are colinear with a genotype which is associated with the phenotype, termed lineage effects (19). Locus effects may be associated with a change in carriage duration due to convergent evolution (which may occur through recombination between lineages). In such regions, the causal loci and corresponding phenotypic effects are easier to identify (35). Linear mixed models can be used to find these variants which are causal for a bacterial phenotype independent of lineage; discovery of homoplasic and polygenic variation associated with the phenotype across the entire tree is well powered (19).

While the high heritability suggests many pathogen variants do affect carriage duration, it does not give information on how many of these will be locus or lineage effects. We mapped carriage duration onto the phylogeny, reconstructing the ancestral state at each node. Consistent with the high heritability of carriage duration we found that carriage length was clearly stratified by lineage (SI figure 4).

We first tested for the association of serotype with carriage duration, which is correlated with sequence type (36) and previously associated with differences in carriage duration (3, 11). We also included resistance to six antibiotics, the causal element to some of which are known to be associated with specific lineages (37). These are therefore possible lineage effects which would be unlikely to be found associated under a model which adjusts for population structure (30).

Not all serotypes and resistances may have an effect on carriage duration, or there may not be enough carriage episodes observed to reach significance. As including extra predictors in a linear regression always increases the variance explained, we first performed variable selection using lasso regression (38) to obtain a more reliable estimate of the amount of variation explained. Where a resistance and serotype are correlated and both associated with a change in carriage duration, this will produce a robust selection of the predictors (39).

The selected predictors and their effect on carriage duration are shown in table 1. The total variance explained by these lineage factors was 0.19, 0.178 for serotype alone and 0.092 for resistance alone. We applied the covariance test (40) to determine which lineage effects significantly affected carriage duration and found that 19F, erythromycin resistance, 23F, 6B caused significant (α < 0.05) increase in carriage duration and being non-typable caused a significant decrease.

**Table 1:** Coefficients from lasso regression model of carriage duration. The mean (intercept) corresponds to a sensitive 6A/C carriage episode, and different serotypes and resistances are perturbations about this mean. Positive effects are expected to have a greater magnitude, due to the positive skew of carriage duration. Rows in bold were significant predictors in the covariance test.

| Factor | Effect on carriage duration (days) |
| --- | --- |
| **Mean (intercept)** | **59.5** |
| **Erythromycin resistance** | **+7.5** |
| Tetracycline resistance | +3.0 |
| Trimethoprim resistance | +2.9 |
| Clindamycin resistance | +1.8 |
| Penicillin intermediate resistance | +1.3 |
| **Serotype 19F** | **+46.9** |
| **Serotype 23F** | **+21.0** |
| **Serotype 6B** | **+16.2** |
| Serotype 14 | +7.2 |
| Serotype 21 | +1.6 |
| Serotype 19B | -0.1 |
| Serotype 18C | -1.9 |
| Serotype 29 | -4.3 |
| Serotype 3 | -4.5 |
| Serotype 4 | -7.2 |
| Serotype 24F | -8.5 |
| **Non-typable (NT)** | **-12.3** |
| Serotype 5 | -18.6 |

We wished to test whether the detected effect of serotype and resistance on carriage duration was entirely mediated through their covariance with lineage, or whether they have an independent and therefore more likely causal relationship. We first looked for differences in duration over three recent capsule gain/loss events; if there is an effect of serotype independent of genetic background, these would be predicted have the largest difference between serotypes while controlling for the relatedness of isolates. No significant difference in duration was seen between isolates with or without capsule within the same lineage (p = 0.39; SI figure 5).

However, as these events were limited in number, assumed genetic independence within the clade and occurred only in part of the population, we also performed the same regression as above while also including lineage (defined by discrete population clusters) as a predictors. This therefore allows serotypes which appear in different population clusters to distinguish whether lineage or serotype had a greater effect on carriage duration. The covariance test found that 19F, erythromycin resistance and non-typable had significant effects on the model (in that order). As these terms enter the model before any lineage specific effect, we conclude that these serotypes and resistances explain some of the narrow-sense heritability.

This suggests that the main lineage effect on carriage duration is the serotype, with a small but significant change caused by whether the bacteria are erythromycin resistant. Our finding that erythromycin resistance significantly affects carriage duration, while being a relatively uncommon treatment in this setting (3% of treatments captured), but that other antibiotics do not may be because erythromycin resistance would be expected to cause an almost four order magnitude increase in MIC, whereas other resistance acquisitions have a much smaller effect.

Additionally, we calculated the mean sojourn times and mean number of carriage episodes from the fit to the HMM for commonly carried serotypes (SI table 2), which gave results similar to the regression performed above. These estimates are comparable to the previous analysis on a subset of these samples. The majority of carriage episodes were due to five of the seven paediatric serotypes (41), or non-typeable isolates. The results show 19F, 23F and 14 were carried the longest, 6A/C and 6B for intermediate lengths, and NT the shortest.

The overall picture of the first two years of infant carriage is one containing one or two long (over 90 day) carriage episodes of a common serotype (6A/C, 6B, 14, 19F, 23F) and around two short (under a month) carriage episodes of non-typable *S. pneumoniae*. Colonisation by other serotypes seem to cause slightly shorter carriage episodes, though the relative rarity of these events naturally limits the confidence in this inference. That some serotypes are rarer and carried for shorter time periods may be evidence of competitive exclusion (42, 43), as fitter serotypes quickly replace less fit serotypes thus leading to reduced carriage duration. The calculated mean carriage duration of NT pneumococci is similar to the minimum resolution we were able to measure by the study design, which suggests carriage episodes may actually be shorter than one month. Unfortunately the only existing study with higher resolution did not check for colonisation by NT pneumococci (3).

These estimates are similar to previous longitudinal studies in different populations (2, 9, 10), though against the Kilifi study our estimates are systematically larger. This may be due to the lower resolution swabbing we performed, or may be because the previous study was unable to resolve multiple carriage (11% of positive swabs). While our heritability estimates are specific to this population due to differences in host, vaccine deployment and transmission dynamics, the similarity of the estimates of serotype effect to those from different study populations suggests our results may be somewhat generalisable.

### Genome wide association study implicates additional locus specific effects in carriage duration

To search for locus effects as discussed above, we applied a linear-mixed model to all the common SNPs and k-mers in the dataset. The results for SNPs are shown in figure 3 and SI table 3, with 14 loci reaching suggestive significance and two reaching genome-wide significance. We also found that 424 k-mers reached genome-wide significance, which we filtered to 321 k-mers over 20 bases long to remove low specificity sequences (SI figure 6). To determine their function, we mapped these k-mers to the coordinates of reference sequences.

**Figure 3:**
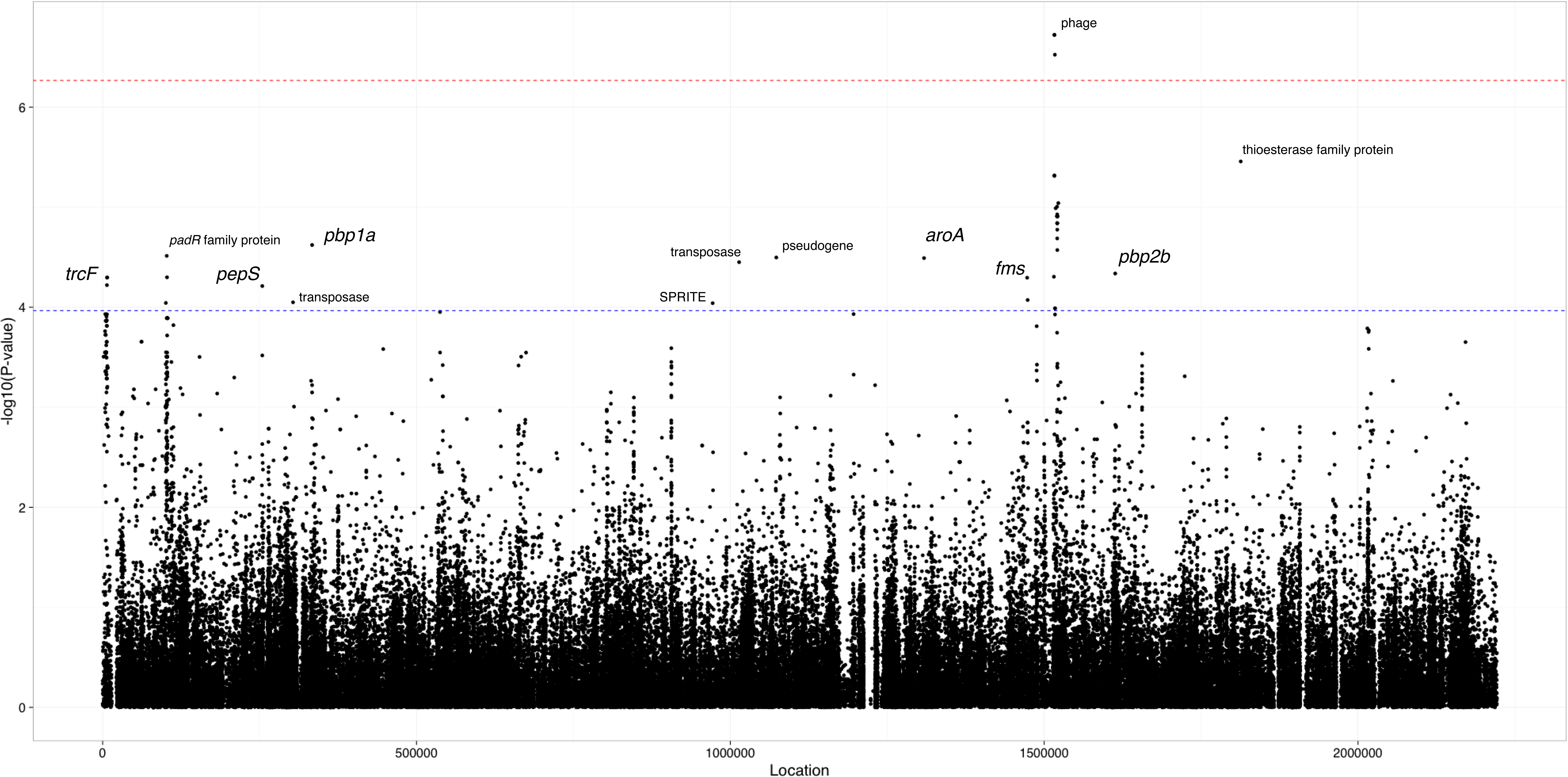
Manhattan plot of SNPs associated with carriage duration. The significance of each SNP’s association with carriage duration against its position in the ATCC 700669 genome is shown. The red line denotes genome-wide significance, and the blue line suggestive significance. Loci reaching suggestive significance are labelled with their nearest annotation, as in SI table 3.

The only genome-wide significant SNP hits are in a prophage in the ATCC 700669 genome (44) (SI figure 7) with the LD structure suggesting there are two separate significant signals found in this region. The most significant k-mer hits were also located in phage sequence and were associated with a reduced duration of carriage. The discovery of associated phage variants have more power under a LMM: the LD block size in this region is smaller than in the rest of the genome, as prophage sequence is highly variable within *S. pneumoniae* lineages (45). Multiple independent phage variants may therefore affect carriage duration, which will increase their significance using a LMM. Indeed, the significant k-mers from the LMM are not significant under a model of association using fixed effects to control for population structure, and are strongly associated with the population structure components of the model.

We postulated that presence of any phage in the genome may cause a reduction in carriage duration. By using the presence of phage as a trait under the linear mixed model, we however found no evidence of association (p = 0.35). These results are therefore evidence that infection with a specific phage sequence is associated with a slight decrease in carriage duration. A similar result has previously been found in a genome-wide screen in *N.meningitidis*, where a specific phage sequence was found to affect the virulence and epidemiology of strains (46, 47).

Signals at the suggestive level include *pbp1a* and *pbp2b*, which suggest as above that penicillin resistance may slightly increase carriage duration, but there are not enough samples in this analysis to confirm or refute this. Other signals near genes at a suggestive level included SNPs in *trcF* (transcription coupled DNA repair), *padR* (repressor of phenolic acid stress response), *pepS* (aminopeptidase), *aroA* (aromatic amino acid synthesis), *fms* (peptide deformylase) and a thioesterase superfamily protein. K-mers from erythromycin resistance genes (*ermB*, *mel*, *mef*) were expected to reach significance from the above analysis, but did not: it has however previously been shown that the power to detect these elements in a larger sample set taken from the same population is limited due to the multiple resistance mechanisms and stratification of resistance with lineage (37).

We found that the test statistic from fast-lmm was underinflated (*λ*_gc_ = 0.84, SI figure 8) suggesting limited power to detect effects associated with both the lineage and phenotype. This effect has been previously noted, and while LMMs have improved power for detecting locus specific effects they lose power when detecting associated variants which segregate with background genotype (19).

To search for candidate regions which may be independently associated with both a lineage and increased carriage duration, we ran an association test with a less stringent population structure correction. This is expected to have higher power than an LMM for true associated variants on ancestral branches, but will also increase the number of false positives (variants co-occurring on these branches which do not directly affect the carriage duration themselves). We observed a highly overinflated test statistic, as expected for this level of population stratification (*λ*_gc_=3.17, SI figure 9).

The most highly associated SNPs were in all three pbp regions associated with *β*-lactam resistance, the capsule locus, *recA* (DNA repair and homologous recombination), *bgaA* (beta-galactosidase), *phoH*-like protein (phosphate starvation-inducible protein), *ftsZ* (cell division protein) and *groEL* (chaperonin). Associated k-mers were also found in *recU* (DNA repair), *phtD* (host cell surface adhesion), *mraY* (cell wall biosythesis), *tlyA* (rRNA methylase), *zinT* (zinc recruitment) and *recJ* (DNA repair). It is not possible to determine whether variation in these genes is associated with a change in carriage duration or if the variation is present in longer carried, generally more prevalent lineages. For example, *β*-lactam resistance may appear associated as the long carried lineages 19F and 23F are more frequently resistant, or it may genuinely provide an advantage in the nasopharynx that extends carriage duration independent of other factors. Future studies of carriage duration, or further experimental evidence will be needed to determine which is the case for these regions.

Antigenic variation in known regions (of *pspA, pspC, zmpA* or *zmpB*) or variation in bacteriocins (*blp*) may both be expected to cause a change in carriage duration (48, 49), however we found neither to be associated with a change in carriage duration. This was likely due to stratification of variation in these regions with lineage, but may also be caused by a larger diversity of k-mers in the region reducing power to detect an association.

### Child age independently affects variance in carriage duration

Finally, we wished to determine the importance of two environmental factors which are known to contribute to variance in this phenotype: child age and whether the carriage episode is the first the child has been exposed to (3, 11, 50). These have been applied throughout the analysis as covariates, both in the estimation of carriage episodes and in associating genetic variation with change in carriage duration.

We applied linear regression to these factors, which showed they were both significantly associated with carriage duration as expected (p<10^−10^). Together, they explained 0.046 of variation in carriage duration. Both remained significant when adjusting for the effect of serotype and drug resistance. As found previously, increasing child age contributes to a decrease in the duration of carriage episodes. From a mean of 68 days long, we calculated a drop of 19 days after a year, and 32 days after two years. Extrapolating, this causes carriage episodes longer than two days to cease by age 11 (SI figure 10). Previous carriage of any serotype was estimated to cause an increase in the duration of future carriage episodes, though previous studies have found no overall effect (51). It has previously been shown that prior exposure to non-typables in this cohort make colonisation by another non-typable occur later, and for a shorter time (11). The positive effect observed in this analysis is therefore likely to be an artefact due to subsequent carriage episodes being more likely to be due to typable pneumococci.

Additional environmental factors that explain some of the remainder of the variance may include the variation of the host immune response and interaction with other infections or co-colonisation. In particular, co-infection influenza A was not recorded but is known to affect population dynamics within the nasopharynx (52). Fundamentally, imprecise inference of the carriage duration will limit our ability to fully explain its variance.

## Discussion

By developing models for longitudinal swab data and combining the results with whole genome sequence data we have quantified and mapped the genetic contribution to the carriage duration of *S. pneumoniae*. We found that despite a range of other factors such as host age which are known to cause carriage duration to differ, sequence variation of the pneumococcal genome explains most of this variability (63%). Common serotypes and resistance to erythromycin cause some of this effect (19% total), as does the presence or absence of particular prophage sequence in the genome. Table 2 summarise the sources we found to be significantly associated with variation in carriage duration.

**Table 2:** Summary of variance of carriage duration explained by genetic and environmental factors. *H*^2^ encompasses all rows, other than the measured environmental effects.

| Source | of which is | Total variance explained | Proportion of total heritability explained |
| --- | --- | --- | --- |
| Total heritability ($H^2$) | | 0.634 | 1.00 |
| | Total common SNP heritability ($h_{SNP}^2$) | 0.438 | 0.691 |
|  | Serotype and resistance | 0.190 | 0.300 |
|  | Serotype only | 0.178 | 0.281 |
|  | Resistance only | 0.092 | 0.145 |
|  | Phage k-mers | 0.067 | 0.106 |
| Measured environmental effects | Age and previous carriage | 0.046 | - |

We provide a quantitative estimate of how closely transmission pairs share their carriage duration, and show evidence for differences both between and within serotypes. The implication of phage as having a significant effect on carriage duration has interesting corollaries on pneumococcal genome diversification through frequent infection and loss of prophage, even during carriage episodes in this dataset.

The results presented here have important implications for the modelling of pneumococcal transmission and their response to perturbation of the population by vaccine. Importantly, our analysis of heritability shows that variants other than serotype affect carriage duration, consistent with recent theoretical work (18). Together these studies suggest that variants exist in the pneumococcal genome which alter carriage duration, which in turn is linked to antibiotic resistance.

We were not able to fully explain the basis for heritability of carriage duration for a number of reasons. The close association of the phenotype with lineage limited our power to fine-map lineage associated variants other than capsule type which may affect carriage duration. Meta-analysis with more large studies with higher resolution may help to resolve these issues: we are conducting a similar study in Cape Town, South Africa will combine sequence data with two-weekly swabs which will be compared to these results in future. Additional environmental factors that explain some of the remainder of the variance may include the variation of the host immune response and interaction with other infections or co-colonisation. In particular, co-infection influenza A was not recorded but is known to affect population dynamics within the nasopharynx (52).

This is a phenotype which would have been difficult to assay by traditional methods such as in an animal model due to the cohort size needed and the length of time experiments would need to be run for. By instead using genome-wide association study methods we have been able to quantitatively investigate a complex phenotype in a natural population. We believe that the analysis of heritability and variance explained in a phenotype of interest, as presented here, will be an important part analysis of complex bacterial traits in future studies.

## Methods

### Sample collection

The study population was a subset of infants from the Maela longitudinal birth cohort (53), and was split into two cohorts. In the ‘routine’ cohort, 364 infants were swabbed monthly from birth, 24 times in total. All swabs were cultured and serotyped using the latex sweep method (20). In the ‘immunology’ cohort 234 infants were swabbed on the same time schedule, but cultured and serotyped following the World Health Organisation (WHO) method (11). Non-typable pneumococci were confirmed by bile solubility, optochin susceptibility and Omniserum Quellung negative. For both cohorts phenotypic drug resistance to six antibiotics was available (chloramphenicol, *β*-lactams, clindamycin, erythromycin, trimethoprim and tetracycline). 3161 randomly selected pneumococcal positive swabs from the study population have been previously sequenced, 2175 of which were from these longitudinal infant samples (33).

### Converting swab data into a time series

Latex sweeps could not differentiate 6A and 6C serotypes, so we treated these as a single serotype when detected by this method (in WHO serotyping PCR was used to differentiate these serotypes). 15B and 15C serotypes spontaneously interconvert, so were combined. We removed two duplicated swabs (08B09098 from the immunology cohort; 09B02164 from the routine observation cohort).

To get a good fit of the HMM, we normalised observation times for each sample. Defining infant birth as *t* = 0, subsequent sampling times *t_i_* were measured in days, and normalised to have a variance of one. The actual (untransformed) carriage duration in days was used as initial phenotype *y*.

### Hidden Markov model of time series

We modelled the time series of swab data using a continuous-time HMM, as implemented in the R package *msm* (54). Unobserved (true) states correspond to whether the child is carrying bacteria in their nasopharynx, and observed (emitted) states correspond to whether a positive swab was seen at each point. Transition probabilities between each state **Q** and the emission probabilities **E** are jointly estimated by maximum likelihood using the BOBYQA algorithm. We then constructed the most likely path through the unobserved states for each child using the Viterbi algorithm (55) with the observed data and estimated model parameters. Assuming that continuous occupation of the carried state corresponded to a single carriage episode, we calculated the duration for each such episode from the inferred true states.

### Processing genetic data

For each isolate with an inferred carriage duration (N = 2175) we extracted SNPs from the previously generated alignment against the ATCC 700669 genome (56). Consequences of SNPs were annotated with VEP, using a manually prepared reference (57). A phylogenetic tree was generated from this alignment using FastTree under the GTR+gamma model (58). The carriage duration was mapped on to this phylogeny using phytools (59). We then filtered the sites in the alignment to remove any where the major allele was an N, any sites with a minor allele frequency lower than 1%, and any sites where over 5% of calls were missing. This left 115210 sites for association testing and narrow-sense heritability estimation.

We counted 68M non-redundant k-mers with lengths 9-100 from the *de novo* assemblies of the genomes using a distributed string mining algorithm (60, 61). We filtered out low frequency variants removing any k-mers with a minor allele frequency below 2%, leaving 17M for association testing.

We identified the presence of phage by performing a blastn of the *de novo* assemblies against a reference database of phage sequence (62). If the length of the top hit was over 5000 we defined the isolate as having phage present (SI figure 11).

### Transformation of carriage duration phenotype

As we aimed to fit a multiple linear regression model to the carriage duration *Y* at each genetic locus *k*, we first ensured the data was appropriate for this model. The phenotype distribution was positively skewed, with an approximately exponential distribution (SI figures 2 and 12). Residuals were therefore non-normally distributed, potentially reducing power (63).

In the regression setting, a monotonic function can be applied to transform the response variable to avoid this problem. We took the natural logarithm of the carriage duration

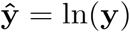

which led to the residuals being much closer to being normally distributed (sup fig X). We applied the same transformation to child age, when it was used as a covariate in association.

### Estimation of heritability

We estimated broad sense heritability *H*^*2*^ with the ANOVA-CPP method in the *patherit* R package (64), using a patristic distance cutoff of 0.04 (SI figure 13).

To estimate the SNP-based heritability *h*^2^_*SNP*_ we applied a linear mixed model, which uses the genomic relatedness matrix (as calculated from SNPs passing filtering) as random effects. We used the implementation in warped-lmm (29), which learns a monotonic transform as it fits the model to the data to ensure residuals are normally distributed (SI figure 3). We therefore used the untransformed phenotype *y* as the input. Child age and whether previous carriage had occurred were included as covariates. We also estimated *h*^2^_*SNP*_ using LDAK (65) with default settings, which gave an estimate of 0.437 (<1% difference from the warped-lmm estimate).

### Association of antimicrobial resistance and serotype with carriage duration

We encoded all 56 observed serotypes (including non-typables) and resistance to the six antibiotics as dummy variables. We used 6A/C as the reference level, as this had a mean carriage duration close to the grand mean in previous analysis. Orthogonal polynomial coding was used for the latter four antibiotics, where resistance could be intermediate or full. We then regressed this design matrix **X** was against the transformed carriage duration *ŷ*. We removed three observations with low carriage lengths due to a delayed initial swab, and seven observations with leverages of one (SI figure 1).

We performed variable selection using lasso regression (38), implemented in the R package *glmnet* (66). We used leave-one-out cross-validation to choose a value for the *ℓ*_1_ penalty; the value one standard error above the minimum cross-validated error (67) was selected (*λ* = 0.033; SI figure 14). The 20 predictors with non-zero coefficients in the model at this value of *λ* (Table 1) were used in a linear regression to calculate the multiple *R*^2^, which corresponds to the proportion of variance explained by these predictors.

Capsule switch events had been previously identified by first reconstructing of the ancestral state of the serotype at each node through maximum parsimony (33). For each node involving loss or gain of the capsule, those with at least one child being a tip were selected to find recent switches (all were capsule gain). The carriage duration of all unencapsulated children of the identified node were used as the null distribution to calculate an empirical p-value for the switched isolate. P-values were combined using Fisher’s method (68).

### Genome wide association of carriage duration

We used the linear mixed model implemented in fast-lmm (69) to associate genetic elements with carriage duration, independent of overall lineage effects. We used the warped phenotype as the response, the kinship matrix (calculated from SNPs) as random effects, and variant presence, child age and previous carriage as fixed effects. For SNPs we used a Bonferroni correction with *α* = 0.05 and an N of 92487 phylogenetically independent sites to derive a genome-wide significance cutoff of P < 5.4×10^−7^, and a suggestive significance cutoff (70) of P = 1.1×10^−4^. Significant SNPs were LD-pruned with a cutoff of R^2^ < 0.2 to define significant loci.

For k-mers we counted 5254876 phylogenetically independent sites, giving a genome wide significance cutoff of 9.5×10^−9^. We used blastn with default settings to map the significant k-mers to seven reference genomes (ATCC 700669, INV104B, OXC141, SPNA45, Taiwan19F, TIGR4 and NT_110_58), and the possible Tn*916* sequences (36).

To search for variants with some level of lineage independence we used SEER (37), using the default of a three dimensional projection of the kinship matrix to control for population structure. We performed association tests on SNPs and k-mers passing the default filters using multiple linear regression, and report the top hits with p < 10^−14^. Significant k-mers were mapped as above.

## Acknowledgements

We would like to thank Susannah Salter for sharing data on non-typable culture positive rates. We also wish to thank Ben Cooper, Doug Speed, and the attendees of the 8th Permafrost workshop (particularly Jukka Corander, Christophe Fraser, Sonja Lehtinen and Johan Pensar) for comments on this work.

Work at the Wellcome Trust Sanger Institute was supported by Wellcome Trust (098051). JAL was supported by a Medical Research Council studentship grant (1365620). NJC was supported by a Sir Henry Dale Fellowship, jointly funded by the Wellcome Trust and the Royal Society (grant number 104169/Z/14/Z). PT was supported by the Wellcome Trust (Grant No. 083735/Z/07/Z). SMRU is part of the Mahidol Oxford Tropical Medicine Research Unit supported by the Wellcome Trust. The funders had no role in study design, data collection and analysis, decision to publish, or preparation of the manuscript.

